# Local adaptation and hybrid failure share a common genetic basis

**DOI:** 10.1101/520809

**Authors:** Greg M. Walter, J. David Aguirre, Melanie J Wilkinson, Thomas J. Richards, Mark W. Blows, Daniel Ortiz-Barrientos

**Author notes:** Corresponding Author: Greg M. Walter, Address: School of Biological Sciences, Life Sciences Building, University of Bristol, Bristol, UK BS8 1TQ.

## Abstract

Testing whether local adaptation and intrinsic reproductive isolation share a genetic basis can reveal important connections between adaptation and speciation. Local adaptation arises as advantageous alleles spread through a population, but whether these same advantageous alleles fail on the genetic backgrounds of other populations remains largely unknown. We used a quantitative genetic breeding design to produce a late generation (F4) recombinant hybrid population by equally mating amongst four contrasting ecotypes of a native Australian daisy for four generations. We tracked fitness across generations and measured morphological traits in the glasshouse, and used a reciprocal transplant to quantify fitness in all four parental habitats. In the glasshouse, plants in the second generation showed a reduction in fitness as a loss of fertility, but this was fully recovered in the following generation. The F4 hybrid lacked extreme phenotypes present in the parental ecotypes, suggesting that genes reducing hybrid fitness were linked to traits specific to each ecotype. In the natural habitats, additive genetic variance for fitness was greatest for habitats that showed stronger native-ecotype advantage, suggesting that a loss of genetic variation present in the parental ecotypes were adaptive in the natural habitats. Reductions in genetic variance for fitness were associated with a loss of ecological trade-offs previously described in the parental ecotypes. Furthermore, natural selection on morphological traits differed amongst the parental habitats, but was not predicted to occur towards the morphology of the parental ecotypes. Together, these results suggest that intrinsic reproductive isolation removed adaptive genetic variation present in the parental ecotypes. The evolution of these distinct ecotypes was likely governed by genetic variation that resulted in both ecological trade-offs and intrinsic reproductive isolation among populations adapted to contrasting environments.

## Introduction

A long-standing goal in evolutionary biology is to quantify how natural selection connects adaptation and speciation. Natural selection acts largely upon the additive effects of genes to promote adaptation (Hill et al. 2008), and species form when incompatible interactions among genes create intrinsic reproductive isolation between diverging lineages (Dobzhansky 1937; Muller 1942). Thus, alleles beneficial in an environment promote adaptation, but whether the same alleles that drive adaptation also create reproductive isolation among taxa remains an open question (Baack et al. 2015). Ultimately, we are interested in understanding how adaptation occurs in different environments and whether speciation and adaptation share a common genetic basis (Coyne and Orr 2004; Funk et al. 2006).

Molecular approaches identifying genes connecting adaptation and speciation are notoriously challenging when applied to complex traits or reproductive isolation (reviewed in Fishman and Sweigart 2018). Complex traits are often polygenic and affected by alleles that differ among individuals in a population. As a consequence, mapping adaptive loci is either technically difficult or biased towards identifying the few segregating loci of large effect (Rockman et al. 2010). Mapping genes underlying reproductive isolation requires directed crosses between taxa that show hybrid sterility or inviability, causing most of the required crosses to fail (Coyne and Orr 2004). These problems are augmented in non-model systems where the lack of genetic resources, and the difficulty in quantifying natural selection in field experiments, challenge our ability to identify whether alleles underlying adaptation are also involved in the evolution of reproductive isolation (Presgraves 2010).

Here, we implement a quantitative genetic approach to explore the connection between adaptation and speciation focusing on estimating additive genetic variation common to both adaptation and reproductive isolation. This approach complements molecular genetic efforts by providing a population-level view of adaptation, and its consequences for segregation of fitness among populations (Demuth and Wade 2005). Specifically, we synthesise late-generation artificial hybrids between locally adapted populations, which we use to explore the consequences of intrinsic reproductive isolation for segregating phenotypic variation, and for genetic variance in fitness in the natural habitats.

In linking adaptation and speciation using hybrids, we must consider the effect of combining divergent genomes on the genetic variation of later hybrid generations. Ideally, we would conserve all genetic variation of the parental taxa, and use recombination to break-up combinations of alleles unique to each parent (Schluter 2000). However, combining divergent genomes can result in a loss of genetic variation when derived alleles are genetically incompatible in alternative genetic backgrounds, creating intrinsic reproductive isolation (Dobzhansky 1937; Muller 1942). We can better connect adaptation and speciation if we can identify the effect of losing ecotype-specific variance (due to intrinsic reproductive isolation) on the segregation of traits in later generation hybrids, and how this might affect responses to natural selection in the natural habitats of the parental populations.

We focus on four recently derived, but contrasting ecotypes of an Australian native wildflower species complex (*Senecio lautus*, Forster f. ex Willd, 1803, *sensu lato*). Ecotypes occur on coastal sand dunes (Dune ecotype) and rocky headlands (Headland ecotype), and in dry sclerophyll woodland (Woodland ecotype) and subtropical rainforest edges (Tableland ecotype) (Ali 1969; Radford et al. 2004; Roda et al. 2013). Previous reciprocal transplant experiments showed that ecotypes in this complex are adapted to their contrasting habitats, and show fitness trade-offs when transplanted into alternative habitats (Walter et al. 2016; Walter et al. 2018b). Genetic variance underlying their contrasting multivariate phenotypes was found to have diverged, and differences in genetic variance were associated with morphological divergence, suggesting that the evolution of genetic variance facilitated adaptive radiation in these ecotypes (Walter et al. 2018a).

Here, we extend this work by connecting patterns of adaptation and speciation in the parental ecotypes by mating among the four ecotypes to create a late-generation F4 hybrid, and testing the consequence of hybridisation in laboratory and field experiments. In creating the F4 generation, we observed a strong reduction in hybrid fertility at the F2 generation, followed by a complete recovery of fitness in the F3 generation (Walter et al. 2016). F2 Hybrid sterility indicated that natural selection assembled beneficial combinations of alleles during the formation of these ecotypes, creating coadapted gene complexes that appear to be easily disrupted during recombination between divergent populations (Fenster et al. 1997; Li et al. 1997; Johansen-Morris and Latta 2006). The fitness recovery in the F3 generation suggests a purge of incompatible ecotype-specific alleles at the F2 generation, and now absent in the F3 generation.

In assessing the consequences of intrinsic reproductive isolation for fitness, we can make an important prediction: If locally adapted alleles that created fitness trade-offs in the wild also created intrinsic reproductive isolation, then alleles involved in local adaptation will be lost after they create genetic incompatibilities that lead to hybrid sterility. As a consequence, future hybrid generations would have reduced genetic variation for fitness in natural habitats, particularly in habitats that heavily favor the native over foreign ecotypes (i.e., show the strongest patterns of local adaptation; Kawecki and Ebert 2004). Moreover, we would also predict that the loss of locally adapted alleles as a consequence of intrinsic reproductive isolation should prevent natural selection on hybrid phenotypes towards the native parental ecotype morphologies. Such tests connect adaptation and speciation by demonstrating that selection for different alleles in different environments leads not only to adaptation, but also the evolution of intrinsic reproductive isolation.

Here, we used a quantitative genetic breeding design to create an F4 hybrid population, which allowed us to estimate additive genetic variance for phenotype in the glasshouse, and for fitness in all four natural habitats. We used extensive field experiments with the F4 recombinant hybrid to understand the ecological significance of genetic variation lost after F2 hybrid sterility and to test the two major hypotheses described above. Our results suggest that genetic variation that is adaptive in different environments creates genetic incompatibilities, and that multivariate phenotypic evolution has important consequences for both adaptation and the evolution of intrinsic reproductive isolation.

## Methods

### Crossing design

To create the F4 we first sampled seeds from one natural population from each of the four ecotypes, which we germinated and grew at the University of Queensland glasshouses. We sampled seeds for the Dune and Headland ecotypes at Lennox Head, NSW (*-28*.*783005, 153*.*594018* and *-28*.*813117, 153*.*605319*, respectively), from the Tableland ecotype at O’Reilley’s Rainforest Retreat, Qld (*-28*.*230508, 153*.*135078*) and the Woodland ecotype at Upper Brookfield, Qld (*-27*.*479946, 152*.*824709*). At each location, we collected seeds from 24-49 plants separated from each other by at least 10m to minimize the likelihood of sampling close relatives. To grow plants, we first scarified each seed and placed them in glass Petri dishes containing moist filter paper. After leaving them in the dark for two days we transferred the germinated seeds to a 25°C constant temperature growth room with 12h:12h light:dark photoperiod. After one week, we transferred the seedlings to the glasshouse and transplanted them into 85mm pots containing a mixture of 70% pine bark and 30% coco peat with slow release osmocote fertilizer and 830g/m^3^ of Suscon Maxi insecticide. We conducted controlled crosses on mature plants by rubbing two mature flower heads together, labeling the flower heads and collecting the seeds as they emerged.

We created the F4 ensuring each ecotype contributed equally and ensuring that at each generation (see Fig 1C), all full-sibling families (hereafter, ‘families’) contributed equally to the next generation. First, we grew plants for the base population from seeds sampled from the natural populations and performed crosses among the ecotypes (*n* = 41-60 individuals/ecotype) to create all combinations of F1 hybrids (*n* = 12 crossing combinations; *n* = 20-25 families/cross type). We then mated among all combinations of crosses in the F1 generation such that all F2 families (*n* = 24 crossing combinations; *n* = 17-22 families /cross type) possessed a grandparent from each of the original parental ecotypes (e.g., F1_Dune,Headland_ × F1_Tableland,Woodland_). Given strong reductions in intrinsic fitness was observed in a previous Dune x Headland F2 hybrid (Walter et al. 2016), we maximized the number of F1 crosses to produce 458 F2 families in total. We grew one individual from each F2 family. Reductions in fitness were observed as F2 hybrid sterility (42% of F2 individuals were successfully mated compared to >90% in F1 hybrids) and reduced fertility (49% reduction in seed set compared to F1 hybrids) (Walter et al. 2016). Consequently, we divided the F2 individuals that produced flowers into three replicate crossing lines to maintain replicates of the construction of the F4. We then randomly mated among all F2 individuals within each line (*n =* 4-12 families/F2 cross type; total F2 families crossed N = 202) to produce the F3 generation (N = 259 families), ensuring that each family contributed equally. We then produced the F4 generation by first growing one individual from each F3 family and randomly designating each individual as a sire or dam. We then mated 115 sires to 114 dams in a full-sibling, half-sibling crossing design to produce 198 families for the F4 generation. The numbers of families and individuals used to create each generation of the F4 are listed in Table S1.

**Fig 1:**
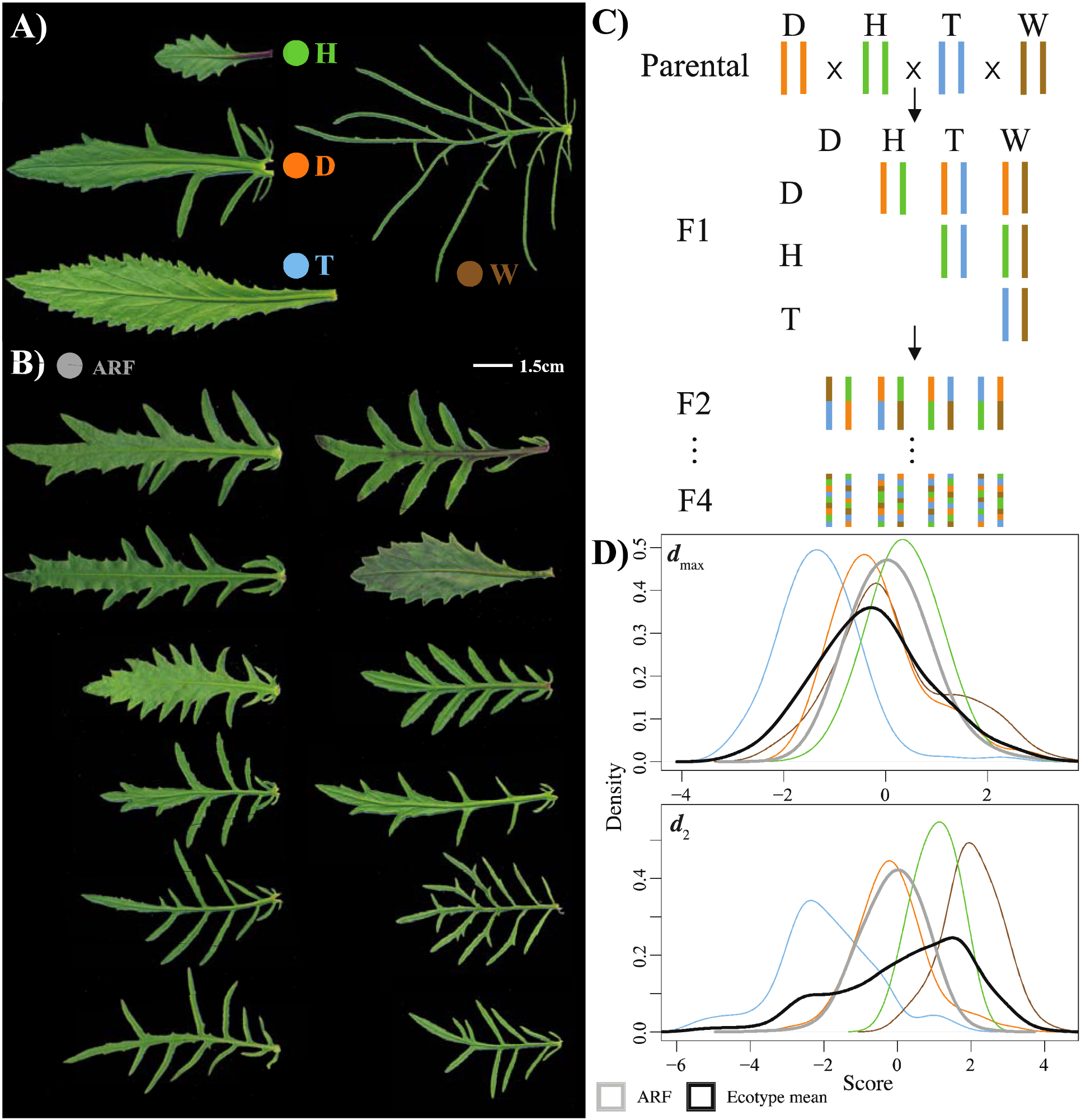
Morphological variation in the ecotypes and F4, alongside a schematic of the crossing design. **A)** Ecotypes vary dramatically in leaf morphology. **B)** The F4 exhibited large variation in leaf morphology, visually intermediate among the original ecotypes. **C)** The F4 was created by equally mating among all ecotypes. **D)** The distribution of ecotype and F4 scores for the first two axes of **D** showing the F4 (grey) occupying an area in phenotypic space similar to the mean of all ecotypes (black) but lacking extreme phenotypes.

We conducted two experiments using the F4. In experiment 1, we grew the F4 in the glasshouse to estimate genetic variance underlying morphological traits. In experiment 2, we transplanted seeds of the F4 into the four habitats to compare the fitness of the F4 with the parental ecotypes. Field fitness of the F4 (experiment 2) was also used to quantify genetic variance in each habitat, and identify genetic-trade-offs among habitats. To quantify differences in natural selection among habitats, we combined both experiments and estimated the genetic covariance between morphological traits in the glasshouse (experiment 1) and field fitness in each habitat (experiment 2).

### Experiment 1: Glasshouse phenotypes

To estimate genetic variance underlying morphological traits we grew four individuals from each full sibling family of the F4 (*n* = 198 full-sibling families, total N = 770 individuals) in 30 cell growth trays containing the same potting media described above. Alongside the F4 we grew four individuals from ~25 full sibling families for each of the parental ecotypes (N = 366 individuals). Plants were grown in a 25°C controlled temperature room with a 12h:12h day:night photoperiod. After eight weeks of growth we measured plant height and sampled one fully mature leaf for each plant. We used the software ‘Lamina’ to analyse the scanned leaf and quantify six variables relating to leaf size and leaf shape (Bylesjo et al. 2008). Using the outputs of Lamina, we quantified leaf morphology using leaf area, leaf area^2^ / leaf perimeter^2^ as a measure of leaf complexity, leaf circularity, number of indents standardized by leaf perimeter, leaf indent width and leaf indent depth.

### Experiment 2: Field transplant

Seeds from the F4 generation of the F4 were transplanted into each of the habitats. At each transplant habitat, we planted 18 seeds from each full-sibling family (*n* = 198) divided equally amongst six experimental blocks (habitat *n* ≈ 3,500 seeds, total N = 14,265 seeds). Alongside the F4 we transplanted seeds from the populations of the parental ecotypes (*n* = 180 seeds/ecotype/habitat) (analysed previously in Walter et al. 2016). See (Walter et al. 2016; Walter et al. 2018b) for a detailed description of the field experiment. Briefly, we glued each seed to a toothpick using non-drip glue and planted them in 25mm×25mm plastic grids in March 2014. To replicate natural germination conditions, we suspended shadecloth (50%) 15cm above each experimental block and watered them daily for three weeks. During the initial three-week period we measured emergence and mortality daily. Following the initial three weeks we measured survival and development at weeks 4, 5, 7 and 9, and then monthly until 20 months at which time there were fewer than 20% of germinated plants remained, and we ceased the experiment. We recorded fitness as: whether each seedling emerged, whether each seedling reached 10 leaves (as a measure of seedling establishment) and produced a bud (reached maturity). All measures of fitness were collected as binary data.

### Implementation of Bayesian models

In the subsequent analyses we implemented Bayesian models to 1) compare field performance of the F4 with the parental ecotypes (experiment 2), 2) identify whether genotype-by-environment interactions create genetic trade-offs among transplant habitats (experiment 2), 3) estimate genetic (co)variance of morphological traits for the F4 (experiment 1), and 4) estimate selection on morphological traits as the genetic covariance between morphological traits (experiment 1) and field performance in each environment (experiment 2). All Bayesian models were implemented using R (R Core Team 2016) within the package ‘MCMCglmm’ (Hadfield 2010). From each model we extracted 1,000 Markov chain Monte Carlo (MCMC) samples, which provided the posterior distribution for the parameters of interest. Details of model specification and convergence checks are located in supplementary material. When estimating genetic variance, comparisons to zero are uninformative. To test whether we estimated biologically meaningful genetic variance, for each Bayesian model implemented on the observed data, we conducted 1,000 randomizations of the raw data to provide a null model for comparisons. Details of how the randomizations were conducted are located in supplementary text.

### Comparing F4 and ecotype morphology

To compare differences in multivariate phenotype between the F4 and parental ecotypes, we implemented a multivariate analysis of variance (MANOVA) on the seven morphological traits measured in experiment 1. We first standardized all seven morphological traits to a mean of zero, and standard deviation of one before including them as a multivariate response variable. To test whether the F4 was phenotypically different to each ecotype we conducted a separate MANOVA for each pairwise comparison between the parental ecotypes, and the F4. We used a bonferroni corrected α-value of 0.0125 (α = 0.05 / *n*, where *n* represents the number of tests). To visualize differences among all ecotypes and the F4 we estimate **D**, the variance-covariance matrix representing multivariate phenotypic divergence. To do so, we first conducted another MANOVA that included all ecotypes (but not the F4). From this, we extracted the sums of squares and cross-product matrices for the ecotypes (SSCP_H_) and error terms (SSCP_E_) to calculate their mean-square matrices by dividing by the appropriate degrees of freedom (MS_H_ = SSCP_H_ / 3; MS_E_ = SSCP_E_ / 365). Using the mean-square matrices we calculated **D** = (MS_H_ – MS_E_) / *nf*, where *nf* represents the number of measured individuals per genotype in an unbalanced design, calculated using equation 9 in (Martin et al. 2008). Our D-matrix then represents divergence in multivariate mean phenotype, among the parental ecotypes, after removing the residual phenotypic variation. To visualize the phenotypic space occupied by the F4 relative to the parental ecotypes, we decomposed **D** into orthogonal axes (eigenvectors) and calculated the scores for the first two eigenvectors for all ecotypes, and the F4.

### (1) Using ecotype-specific variation to connect adaptation and speciation

We estimated fitness at early life history stages for the F4 and parental ecotypes transplanted into all four habitats. We used MCMCglmm to implement the model,

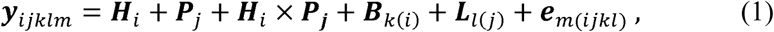

where transplant habitat (***H***_*i*_), F4/ecotype (***P***_*j*_) and their interaction (***H***_*i*_ × ***P***_*j*_) were included as fixed effects. Blocks within transplant habitat (***B***_*k*(*i*)_) and replicate genetic lines within the F4 (***L***_*l*(*j*)_) were included as random effects, and ***e***_*m*(*ijkl*)_ represented the model error. We implemented equation 1 with emergence, seedling establishment and maturity as a multivariate response variable (***y***_*ijklm*_).

We used the F4 to investigate differences in natural selection among contrasting natural habitats. We estimated genetic variance in fitness for each environment, and genetic covariance among environments by quantifying genotype-by-environment (G×E) interactions using a character state approach, where different environments represent different traits. We used the field performance of the F4 and implemented

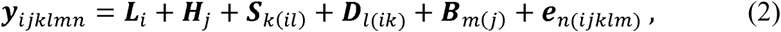

where replicate genetic line of the F4 (***L***_*i*_) and transplant habitat (***H***_*j*_) were included as fixed effects. We included sire (***S***_*k*(*il*)_), dam (***D***_*l*(*ik*)_) and block within habitat (***B***_*m*(*j*)_) as random effects, with ***e***_*n*(*ijklm*)_ representing the residual error variance. For each term in the random component, we estimated random intercepts for each habitat and the covariance among habitats. For the sire and dam components we estimated a 4×4 covariance matrix representing variance in each habitat, and covariance among habitats. Information for estimating covariance among habitats is taken from individuals of the same full-sibling families transplanted in each habitat. Consequently, we implemented equation 2 with a heterogeneous residual covariance matrix, allowing for different variances in each habitat, but fixed residual covariances at zero because individuals (seeds) could not be planted in two habitats simultaneously. We used three separate implementations of equation 2 for emergence, seedling establishment and maturity included as binary univariate response variables (**y**_*ijklmn*_). We extracted the sire variance component from equation 2 (**S**_*l(im)*_) and multiplied by four to estimate the additive genetic (co)variance for field performance among the four habitats.

### (2) Selection on additive genetic variation of complex traits

By linking genetic variance underlying morphological traits in the laboratory, with genetic variance underlying field performance, we sought to quantify differences in natural selection among the transplant habitats using the Robertson-Price Identity. A requirement for natural selection is genetic variance in both morphological traits and field performance. To identify the morphological traits with substantial genetic variation we estimated genetic variance for all traits measured in experiment 1. To estimate genetic variance, we scaled the traits to a mean of zero and standard deviation of one, and implemented

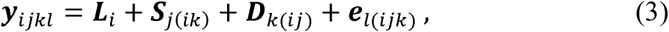

where replicate genetic (***L***_*i*_) was the only fixed effect, with sire (***S***_*k*(*il*)_) and dam (***D***_*l*(*ik*)_) the random effects, and ***e***_*l*(*ijk*)_ representing the residual error. Seven morphological traits were included as a multivariate continuous response (***y***_*ijkl*_). Multiplying the sire variance component by four estimated trait heritabilities, and we found only four traits with heritabilities greater than 0.1 (Table S3; plant height, leaf area, leaf perimeter^2^ / area^2^ and leaf indent width), which we then combined with field performance to study natural selection in the subsequent analyses, described below.

We estimated the genetic covariance between the morphological traits and field performance by implementing the Robertson-Price Identity, R = *s*_*g*_ = *cov*_*a*_ (*w,z*) where the response to selection (R) is analogous to the selection differential (*s*_*g*_), calculated as the additive genetic covariance between a trait (*z*) and fitness (*w*). R generalizes to multivariate form by including more phenotypic traits and estimating a genetic variance-covariance matrix (**G**), with fitness as the final trait. In this framework, *s*_g_ generalizes to the vector of selection gradients (***s***_g_) representing the multivariate response to selection. Estimating the response to selection in this way includes both direct and indirect selection. To estimate selection in the absence of genetic correlations among traits, we calculated the genetic selection gradient (***β***g) using ***β***g = ***G***^**–1**^ ***s***g (Lande and Arnold 1983).

To implement the Roberston-Price Identity we estimated the (co)variance between the four morphology traits and field performance with the linear mixed model

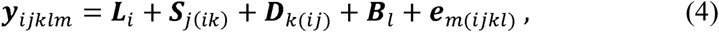

where replicate genetic line (***L***_*i*_) was the only fixed effect. Sire (***S***_*j*(*ik*)_) and dam (***D***_*k*(*ij*)_) were included as random effects along with block within habitat (***B***_*l*_). The multivariate response variable (***y***_*ijklm*_) included four phenotypic traits and the ability to reach maturity in each habitat. Fitness and morphology was measured on separate individuals (field versus glasshouse experiments), and so similar to equation 2, we estimated a heterogeneous residual covariance matrix.

Multiplying the sire variance component (***S***_*j*(*ik*)_) by four (from equation 4) gave the 5×5 additive genetic variance-covariance matrix (**G**). Elements in the first four rows and columns represented **G** among morphological traits. Covariance elements in the fifth column (and row) denote the genetic covariance between each trait and fitness (***s***_**g**_), with genetic variance in fitness in the final element (fifth row, fifth column) (Stinchcombe et al. 2014). We implemented equation 4 separately for field performance in each of the four transplant habitats. We extracted ***s***_**g**_ as the vector of covariances between morphological traits and field performance (rows one to four of the fifth column).

To identify whether we captured biologically meaningful differences in selection among habitats, we conducted two analyses. First, ***s***_g_ and ***β***_g_ being vectors, if the observed length of the vectors (dot product) were greater than the length calculated from the randomised matrices, there is evidence we detected biologically meaningful estimates of selection (Stinchcombe et al. 2014; Walsh and Lynch 2018). Second, we quantified differences in ***s***_g_ among habitats using

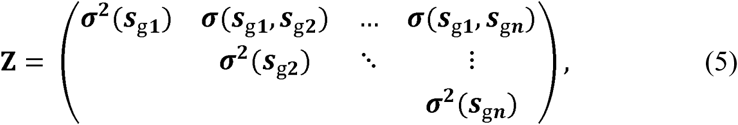

where **Z** represents the among-habitat variance in ***s***_**g**_ for the *n*th trait along the diagonal. The off-diagonal then contains the covariance in ***s***_g_ among habitats, for each bivariate trait combination (Chenoweth et al. 2010). In the same way, we used equation 5 to calculate **B**, the among-habitat (co)variance in ***β***_g_. Comparing the eigenvalues from the observed and random matrices (for both **B** and **Z**) tested whether we captured biologically meaningful differences in selection.

## Results

### Comparing F4 and ecotype leaf morphology

The F4 generation hybrid (F4) was created by equally combining plants from populations of all four ecotypes (Dune, Headland, Tableland and Woodland; Fig 1C). Ecotypes show strong differences in leaf morphology (Fig 1A), with the F4 exhibiting a phenotype intermediate to the parental ecotypes (Fig 1B). Pairwise MANOVAs showed the F4 leaf phenotype was significantly different to all parental ecotypes (Dune: Wilks’ λ = 0.71, *F*_1,857_ = 50.782, *P* = <0.001; Headland: Wilks’ λ = 0.64, *F*_1,863_ = 69.747, *P* = <0.001; Tableland: Wilks’ λ = 0.38, *F*_1,866_ = 192.36, *P* = <0.001; Woodland: Wilks’ λ = 0.22, *F*_1,864_ = 445.1, *P* = <0.001). The MANOVA conducted on the parental ecotypes described a significant difference in multivariate mean phenotype (Wilks’ λ = 0.03, *F*_3,362_ = 117.86, *P* = <0.001), where differences among ecotypes captured 64% of the total variance. We used the MANOVA to calculate **D**, the covariance matrix representing among ecotype divergence in mean phenotype. The first two axes of **D** (***d***_max_ and ***d***_2_) described 84% and 14% of phenotypic divergence, respectively. ***d***_max_ captured differences between the Tableland and Headland ecotypes, with ***d***_2_ capturing differences between the Woodland and Tableland ecotypes (Fig 1D). The F4 occupied an area in phenotypic space close to the Dune ecotype, and intermediate between the Headland, Tableland and Woodland ecotypes. The F4 mean was similar to the overall mean of all ecotypes, but exhibited lower phenotypic variance, lacking in extreme phenotypes of the Headland, Tableland and Woodland ecotypes and suggesting ecotype specific genetic and phenotypic variation was lost following F2 hybrid sterility (Fig 1D).

### (1) Using ecotype-specific variation to connect adaptation and speciation

As reported previously, native ecotypes performed better than foreign ecotypes in all four transplant environments, especially for seedling establishment in the tableland and headland habitats (Walter et al. 2016). We found the performance of the F4 was similar or higher to the native ecotypes for seedling establishment and maturity (Fig 2), suggesting heterosis after several generations of recombination.

**Fig 2:**
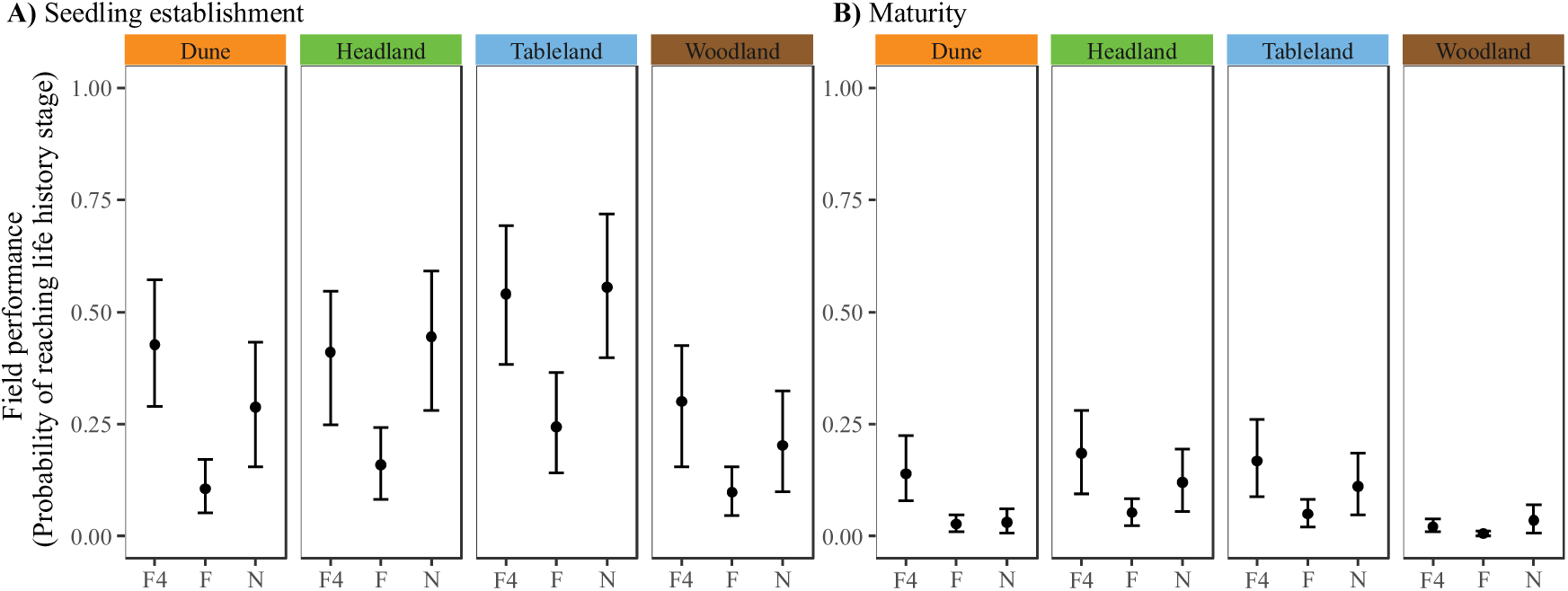
Field performance of the F4 compared to foreign (F) and native (N) ecotypes in each transplant habitat. Fitness measured as the probability of reaching **A)** seedling establishment, and **B)** maturity. Credible intervals represent 95% HPD intervals.

If F2 hybrid sterility removed adaptive alleles from the parental ecotypes, we predicted that as a consequence of losing adaptive alleles the transplant environments that showed strong native-ecotype advantage in Fig 2 would also show reduced additive genetic variance in F4 fitness, and weak genetic trade-offs with the other habitats. To test this hypothesis, we estimated **G**_F4_, the (co)variance matrix representing genetic variance in F4 fitness in each habitat and the genetic covariance among habitats. This matrix describes the amount of genetic variation in fitness for each habitat and quantifies how genetic variation in fitness is shared among habitats. Additive genetic variance for field fitness increased as life-history stages progressed for all habitats (Fig 3). Consistent with our prediction, the habitats associated with stronger native-ecotype advantage (Fig 2, the Headland and Tableland habitats) also showed the least additive genetic variance for fitness in the F4 population for both seedling establishment and maturity (Fig 3).

**Fig 3:**
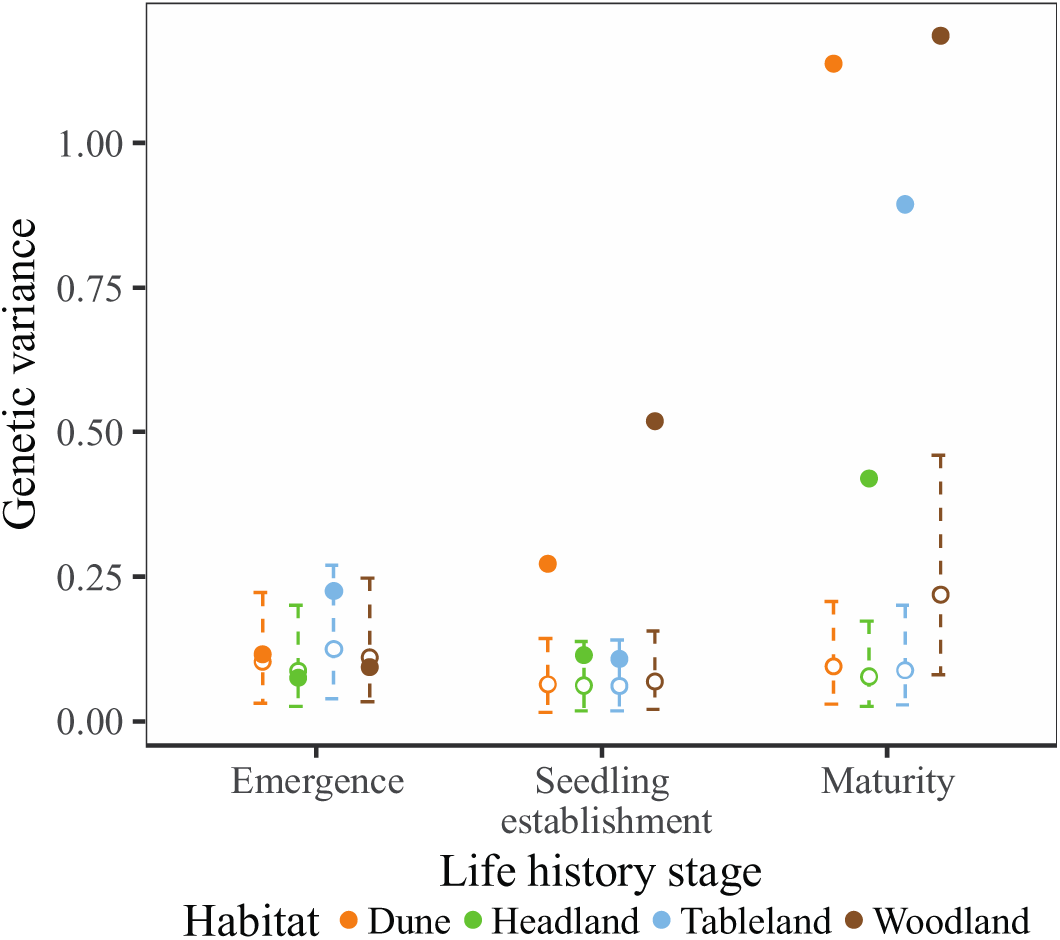
Genetic variance for field performance in the F4 for each habitat (coloured), at each life history stage. Filled circles represent observed estimates, with dashed lines and unfilled circles representing the variance expected by random sampling of fitness values from the population. Credible intervals represent 95% HPD intervals.

To test whether the loss of adaptive alleles (at the F2 generation) reduced the amount of genetic variance in field fitness, we tested for a significant negative correlation between the strength of adaptive divergence in each habitat (Fig 2; native-ecotype performance – foreign-ecotype performance) and the amount of genetic variance in the F4 generation. As predicted, we found a strong negative association for maturity and a weaker negative association for seedling establishment (Fig 4), suggesting alleles associated with adaptive divergence created genetic incompatibilities that reduced genetic variance in F4 fitness. Regardless of the life-history stage, stronger patterns of adaptive divergence were associated with less additive genetic variance for field performance (Fig 4).

**Fig 4:**
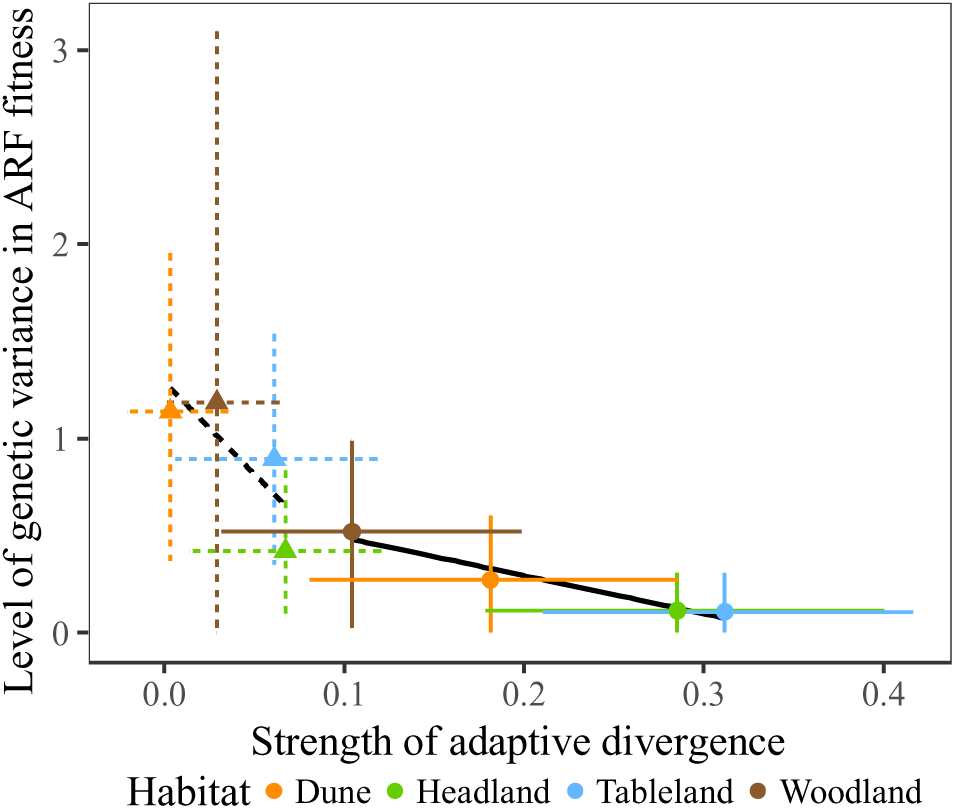
Stronger patterns of adaptive divergence were negatively associated with the level of genetic variance. Strength of adaptive divergence was quantified as the difference in fitness between the native and foreign ecotypes. Solid circles and lines represent seedling establishment, triangles and dashed lines represent maturity. Credible intervals represent 95% HPD intervals. Estimating separate regression slopes for each MCMC iteration showed that the distribution of the slope was negative and did not overlap zero at 88% HPD for maturity but overlapped zero completely for seedling establishment.

Decomposing **G**_F4_ showed that the first three axes described more genetic variance in fitness than expected by random sampling (Fig S1). The first axis of among-habitat genetic variance in fitness described a common genetic basis shared among all habitats, which suggested it described heterosis or genetic variation needed to function across all four transplant environments (Table 1). Axis two and axis three of **G**_F4_ provided evidence of genetic tradeoffs, describing genetic variance in fitness that differed between the woodland and dune ecotypes (***e***_2_), as well as between the tableland, and the dune and woodland transplant habitats (***e***_3_). Therefore, in addition to a common genetic basis to fitness in all environments (***e***_1_; Table 1) we detected genetic trade-offs for fitness among habitats (Table 1). Notably, the tableland and headland habitats showed the highest levels of native-ecotype advantage in Fig 2 but were weakly associated with F4 fitness trade-offs. The matrices for all life history stages are presented in supplementary Table S2.

**Table 1:** Eigenanalysis of the additive genetic (co)variance matrix for field performance at maturity. Loadings in bold are greater than 0.25 to aid interpretation. HPD represents the observed 95% HPD credible intervals.

| Habitat | Eigenvectors | $e_1$ | $e_2$ | $e_3$ | $e_4$ |
| --- | --- | --- | --- | --- | --- |
|  | Eigenvalue | 2.492 | 0.782 | 0.248 | 0.116 |
|  | HPD | 0.837-4.179 | 0.036-1.984 | 0.037-0.569 | 0.011-0.267 |
|  | Proportion | 0.685 | 0.215 | 0.068 | 0.032 |
|  | Dune | **-0.56** | **0.61** | **0.45** | **-0.34** |
|  | Headland | **-0.34** | 0.22 | -0.04 | **0.91** |
|  | Tableland | **-0.54** | -0.02 | **-0.81** | **-0.23** |
|  | Woodland | **-0.53** | **-0.76** | **0.37** | 0.00 |

### (2) Selection on additive genetic variation of complex traits

To quantify selection in each habitat we calculated the multivariate response to selection (***s***_g_) as the additive genetic covariance between morphological traits measured in the glasshouse, and performance measured in the field for each of the four habitats (the Robertson-Price Identity; Robertson 1966; Price 1970). To estimate selection in the absence of genetic correlations we calculated ***β***_g_, the genetic selection gradient for each habitat. Comparing the observed and random estimates of ***s***_g_ and ***β***_g_ provided evidence that we captured biologically meaningful selection within each habitat, and meaningful differences in selection among habitats (Fig S2). We then quantified differences among habitats in both ***s***_g_ and ***β***_g_. To do so, we estimated **Z** and **B**, the covariance matrices representing the among-habitat difference in ***s***_g_ and ***β***_g_, respectively.

We tested whether differences in the response to selection (**Z**) or differences in selection gradients (**B**) predicted evolution towards parental ecotype morphology (**D**). To do so, we decomposed **Z** and **B**, where the first axis of each matrix described 83% (HPD 56-98%) and 81% (HPD 55-98%) of the among-habitat variance, respectively. We predicted that if natural selection occurred in the direction of parental ecotype morphology, differences in selection on the F4 (**Z** and **B**) would capture more variation in **D** than expected by chance. This is important because if we detect selection towards the native-ecotype morphology, then genetic incompatibilities had no influence on adaptive genetic variation underlying divergence in morphological traits. We found that although selection on the F4 differed among habitats, it did not occur in directions similar to the parental ecotypes (Fig 5A). Consequently, we found that neither the response to selection (first axis of **Z**) nor differences in selection gradients (first axis of **B**) aligned with the first axis of **D, *d***_max_ (Fig 5B). However, the greatest differences in selection aligned with ***d***_2_, the second axis of phenotypic divergence in the parental ecotypes (Fig 5C).

**Fig 5:**
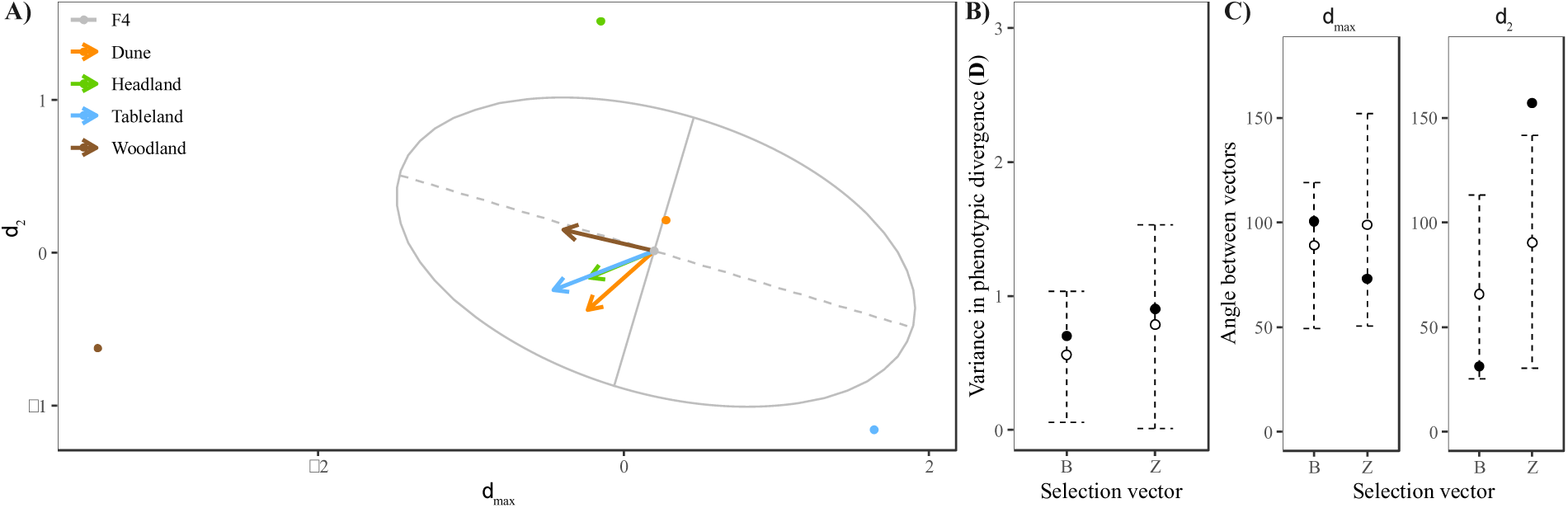
Selection on the F4 did not predict evolutionary trajectories. **A)** Visualising selection on the F4 by calculating the F4 G-matrix (ellipse) and selection gradients (arrows) within the space of the D-matrix, phenotypic divergence of the parental ecotypes. **B)** Differences in Z and B did not describe significant divergence in parental morphology. **C)** Differences in selection aligned with closer with ***d***_2_, but not ***d***_max_.

## Discussion

While there is abundant evidence implicating divergent natural selection in the accumulation of extrinsic reproductive barriers such as immigrant inviability and ecologically dependent postzygotic isolation (Rundle and Nosil 2005), the contribution of local adaptation to the evolution of genetic incompatibilities during population divergence remains unresolved (Baack et al. 2015). Many genes underlying postzygotic isolation show the signature of past rapid evolution, but studies connecting genes underlying both local adaptation and reproductive isolation are rare (Presgraves 2010). Our results clarify the connection between adaptation and the evolution of intrinsic reproductive isolation by showing that F2 hybrid sterility can be associated with the loss of extreme phenotypic variation, and that such alleles likely contributed to adaptive ecotypic divergence. Our work also shows that future research should consider the evolution of genetic incompatibilities in a quantitative genetic framework to understand how additive genetic variation is lost due to intrinsic reproductive isolation.

### The nature of coadapted gene complexes

F2 hybrid sterility followed by fitness recovery in the subsequent F3 generation indicates population divergence as a build-up of coadapted gene complexes (Rawson and Burton 2002; Johansen-Morris and Latta 2006). Coadapted gene complexes can be considered in two ways. First, they can be epistatic interactions between genes that work well within a species, but that fail in the background of related species (Dobzhansky 1937; Muller 1942). In this case, positive epistatic interactions are required to function, and recombination (regardless of the genetic makeup of the other taxa) shuffles alleles, which creates negative epistasis that results in intrinsic hybrid sterility or inviability. Coadapted gene complexes can often evolve rapidly (e.g., Presgraves and Stephan 2007), and are known to produce genetic incompatibilities causing hybrid inviability and sterility (Brideau et al. 2006; Tang and Presgraves 2009; Bomblies 2010; Nosil and Schluter 2011). Alternatively, coadapted gene complexes can be viewed as a build-up of beneficial combinations of alleles maintained by natural selection whose additive effects promote adaptation of populations to local environments (Johansen-Morris and Latta 2006). Intrinsic reproductive isolation would then arise when derived alleles from different populations are combined, creating genetic combinations that reduce hybrid fitness in all environments. In this scenario, extrinsic postzygotic reproductive isolation (selection against hybrids) will evolve when derived alleles from different populations are combined, creating intermediate additive effects that reduce hybrid fitness in the habitat of their parents (Rundle and Whitlock 2001).

Our results suggest that the genetic basis of intrinsic and extrinsic hybrid failure might have a common basis where some of the additive effects contributing to local adaptation and extrinsic postzygotic isolation are also part of epistatic interactions that fail in hybrids. Because intrinsic postzygotic isolation occurs in any environment, but extrinsic reproductive isolation only in the natural environments of populations, our data reveals that F2 hybrid sterility removed some alleles that act additively within populations, but negatively interact between populations. The removal of such additive genetic effects alters the distribution of phenotypes in the next generation of hybrids, which reduces the response to selection in the wild.

The study of quantitative genetic variation can also help us understand how genetic incompatibilities arise among populations. For instance, recent work on the genetics of intrinsic reproductive isolation suggests that derived alleles responsible for genetic incompatibilities are not always fixed in different populations and are segregating as polymorphisms in natural populations (Cutter 2012; Sweigart and Flagel 2015; Larson et al. 2018). Creating hybrid populations and tracking additive variance for both fitness and traits can help us survey polymorphic effects of adaptation on reproductive isolation. Estimating additive genetic variance across generations can also quantify whether different population pairs of the same ecotypes (e.g., the Dune and Headland parapatric pairs) share similar additive genetic variance and whether such sets of alleles are important for adaptation and reproductive isolation across a study system. This kind of information can reveal the tempo and mode of speciation, perhaps hinting rapid evolution of reproductive isolation if alleles that promote adaptation also contribute to hybrid sterility even during the early stages of adaptation when they still segregate in a population.

Our experiments also reveal an important genetic architecture of speciation and adaptation: dominant alleles contribute to adaptation within species, but they also lead to negative interactions between ecotypes resulting in intrinsic reproductive isolation (Li et al. 1997; Demuth and Wade 2005). When we created the F2 hybrids for the current study, we mated unrelated F1 hybrids (e.g., F1_Dune,Headland_×F1_Tableland,Woodland_), which increased the heterozygosity of our F2 hybrids compared to traditional F2 hybrids created by mating two parental taxa. F2 sterility was severe, suggesting that dominant interactions were largely responsible for genetic incompatibilities that might otherwise be absent in traditional F2 hybrid populations. It is well known that dominant alleles, compared to recessive alleles, escape demographic sinks when they are rare, and are more likely to be favoured by natural selection (Haldane 1924). Thus, we conjecture that the fixation of dominant alleles concomitantly promotes adaptation and intrinsic reproductive isolation when fitness is reduced as a consequence of dominant alleles failing on other genetic backgrounds (Li et al. 1997; Demuth and Wade 2005).

### Natural selection on additive genetic variance in hybrids

Field experiments, including a study on coastal ecotypes of *S. lautus*, have shown that natural selection on F2 hybrids occurred in the direction of the native phenotypes (Nagy 1997; Richards et al. 2019). This suggests that our inability to detect selection towards native phenotypes in the F4 population is due to the removal of alleles underlying adaptive phenotypes following hybrid failure. The loss of alleles underlying adaptation (following F2 hybrid sterility) will likely alter the distribution of additive genetic variance underlying multivariate phenotypes in the hybrid population. For any particular habitat, lacking the genetic variation upon which natural selection relies will likely constrain the ability of hybrids to move towards the optimal phenotype. Heterosis could also mask differences in selection between habitats (as shown by the shared genetic variation for fitness), but the removal of additive genetic variance at F2 hybrid sterility in fitness suggests that genetic incompatibilities are likely responsible for the missing genetic variation that prevented selection towards the native parental ecotypes.

Whether the new combinations of alleles in hybrids created by recombination among ecotypes will facilitate the colonization of other habitats remains unknown. However, we anticipate that hybrid species could form when negative genetic interactions removed alleles underlying adaptation to specific environments, creating novel genetic variation for hybrid species to adapt to an environment unoccupied by the parental taxa. Hybrids would contain phenotypic variation absent in their parents, allowing them to evolve along different lines of phenotypic evolution, and at the same time be reproductively isolated from their parents. The repeated and independent of origin of hybrid species could also follow, as perhaps seen in African cichlids (Meier et al. 2017). Therefore, our results also have implications for the way we think of the origin of hybrid species (Abbott et al. 2013), and how the mechanisms stabilizing homoploid genomes (Rieseberg 1997) might contribute to the multivariate genetic constraint left by hybridization between divergent ecotypes.

### Conclusion

Our work indicates that the alleles responsible for the negative genetic interactions (that create genetic incompatibilities) are likely members of co-adapted gene complexes that mainly contribute to local adaptation. In other words, these alleles segregate in the additive genetic variance within an ecotype, forming genetic positive or neutral interactions within the genome of an ecotype, but reduce fitness when combined with the genomes of other ecotypes. We can then view the evolution of ecotypes from a perspective where selection acts upon additive genetic variation by concomitantly increasing allele frequencies at independent loci (Hill et al. 2008), that can be maintained by the evolution of limited recombination amongst loci in response to small population sizes, extensive maladaptive gene flow or very strong selection for integrated phenotypes (Mayr 1954; Carson and Templeton 1984; Ortiz-Barrientos et al. 2016). Some of the environment-specific alleles that segregate within an ecotype fail in alternative genetic backgrounds of other ecotypes, creating intrinsic reproductive isolation and speciation.

## Acknowledgements

We are indebted to Mal Smith, Ruth and Andrew Shaw, and the Ballina Shire Council for generously allowing access to the transplant sites. We are very grateful to M. James, C. Palmer, S. Edgley, J. Walter, L. Ambrose, B. Ayalon, S. Carrol, M. Gallo, F. Roda, A. Maynard, and H. North for their help with the fieldwork. This project was funded by Australian Research Council grants to DO-B. The authors declare no conflict of interest.

## Notes

#### Summary of Updates

Title change and general revision of the writing

## References

Abbott, R., D. Albach, S. Ansell, J. W. Arntzen, S. J. E. Baird, N. Bierne, J. W. Boughman et al. 2013. Hybridization and speciation. Journal of Evolutionary Biology 26:229–246.

Ali, S. I. 1969. *Senecio lautus* Complex in Australia .5. Taxonomic interpretations. Australian Journal of Botany 17:161–176.

Baack, E., M. C. Melo, L. H. Rieseberg, and D. Ortiz-Barrientos. 2015. The origins of reproductive isolation in plants. New Phytologist.

Bomblies, K. 2010. Doomed Lovers: Mechanisms of Isolation and Incompatibility in Plants. Annual Review of Plant Biology 61:109–124.

Brideau, N. J., H. A. Flores, J. Wang, S. Maheshwari, X. Wang, and D. A. Barbash. 2006. Two Dobzhansky-Muller genes interact to cause hybrid lethality in *Drosophila*. Science 314:1292–1295.

Bylesjo, M., V. Segura, R. Y. Soolanayakanahally, A. M. Rae, J. Trygg, P. Gustafsson, S. Jansson et al. 2008. LAMINA: a tool for rapid quantification of leaf size and shape parameters. BMC Plant Biology 8.

Carson, H. L., and A. R. Templeton. 1984. Genetic Revolutions in Relation to Speciation Phenomena - the Founding of New Populations. Annual Review of Ecology and Systematics 15:97–131.

Chenoweth, S. F., H. D. Rundle, and M. W. Blows. 2010. The contribution of selection and genetic constraints to phenotypic divergence. The American Naturalist 175:186–196.

Coyne, J. A., and H. A. Orr. 2004, Speciation. Sunderland, MA, Sinauer Associates.

Cutter, A. D. 2012. The polymorphic prelude to Bateson-Dobzhansky-Muller incompatibilities. Trends in Ecology & Evolution 27:209–218.

Demuth, J. P., and M. J. Wade. 2005. On the theoretical and empirical framework for studying genetic interactions within and among species. The American Naturalist 165:524–536.

Dobzhansky, T. G. 1937, Genetics and the origin of species. New York, Columbia University Press.

Fenster, C. B., L. F. Galloway, and L. Chao. 1997. Epistasis and its consequences for the evolution of natural populations. Trends in Ecology & Evolution 12:282–286.

Fishman, L., and A. L. Sweigart. 2018. When Two Rights Make a Wrong: The Evolutionary Genetics of Plant Hybrid Incompatibilities. Annual Review of Plant Biology 69:17.11–17.25.

Funk, D. J., P. Nosil, and W. J. Etges. 2006. Ecological divergence exhibits consistently positive associations with reproductive isolation across disparate taxa. Proceedings of the National Academy of Sciences of the United States of America 103:3209–3213.

Hadfield, J. D. 2010. MCMC Methods for Multi-Response Generalized Linear Mixed Models: The MCMCglmm R Package. Journal of Statistical Software 33:1–22.

Haldane, J. B. S. 1924. A mathematical theory of natural and artificial selection, Part I. Transactions of the Cambridge Philosophical Society 23:19–41.

Hill, W. G., M. E. Goddard, and P. M. Visscher. 2008. Data and theory point to mainly additive genetic variance for complex traits. PLoS Genet 4:e1000008.

Johansen-Morris, A. D., and R. G. Latta. 2006. Fitness consequences of hybridization between ecotypes of *Avena barbata*: Hybrid breakdown, hybrid vigor, and transgressive segregation. Evolution 60:1585–1595.

Kawecki, T. J., and D. Ebert. 2004. Conceptual issues in local adaptation. Ecology Letters 7:1225–1241.

Lande, R., and S. J. Arnold. 1983. The Measurement of Selection on Correlated Characters. Evolution 37:1210–1226.

Larson, E. L., D. Vanderpool, B. A. J. Sarver, C. Callahan, S. Keeble, L. L. Provencio, M. D. Kessler et al. 2018. The Evolution of Polymorphic Hybrid Incompatibilities in House Mice. Genetics 209:845–859.

Li, Z. K., S. R. M. Pinson, A. H. Paterson, W. D. Park, and J. W. Stansel. 1997. Genetics of hybrid sterility and hybrid breakdown in an intersubspecific rice (*Oryza sativa* L.) population. Genetics 145:1139–1148.

Martin, G., E. Chapuis, and J. Goudet. 2008. Multivariate Q_st_–F_st_ Comparisons: A Neutrality Test for the Evolution of the G Matrix in Structured Populations. Genetics 180:2135–2149.

Mayr, E. 1954. Change of genetic environment and evolution, Pages 157–180 in J. Huxley, A. C. Hardy, and E. B. Ford, eds. Evolution as a process. London, Allen and Unwin.

Meier, J. I., D. A. Marques, S. Mwaiko, C. E. Wagner, L. Excoffier, and O. Seehausen. 2017. Ancient hybridization fuels rapid cichlid fish adaptive radiations. Nature Communications 8.

Muller, H. J. 1942. Isolating mechanisms, evolution and temperature. Biology Symposium 6.

Nagy, E. S. 1997. Selection for native characters in hybrids between two locally adapted plant subspecies. Evolution 51:1469–1480.

Nosil, P., and D. Schluter. 2011. The genes underlying the process of speciation. Trends in Ecology & Evolution 26:160–167.

Ortiz-Barrientos, D., J. Engelstädter, and L. H. Rieseberg. 2016. Recombination Rate Evolution and the Origin of Species. Trends in Ecology & Evolution 31:226–236.

Presgraves, D. C. 2010. Darwin and the Origin of Interspecific Genetic Incompatibilities. The American Naturalist 176:S45–S60.

Presgraves, D. C., and W. Stephan. 2007. Pervasive adaptive evolution among interactors of the *Drosophila* hybrid inviability gene, Nup96. Molecular Biology and Evolution 24:306–314.

Price, G. R. 1970. Selection and Covariance. Nature 227:520–521.

R Core Team. 2016. R: A language and environment for statistical computing, version v. 3.3.2. R Foundation for Statistical Computing, Vienna, Austria.

Radford, I. J., R. D. Cousens, and P. W. Michael. 2004. Morphological and genetic variation in the *Senecio pinnatifolius* complex: are variants worthy of taxonomic recognition? Australian Systematic Botany 17:29–48.

Rawson, P. D., and R. S. Burton. 2002. Functional coadaptation between cytochrome c and cytochrome c oxidase within allopatric populations of a marine copepod. Proceedings of the National Academy of Sciences of the United States of America 99:12955–12958.

Richards, T. J., D. Ortiz-Barrientos, and K. McGuigan. 2019. Natural selection drives leaf divergence in experimental populations of *Senecio lautus* under natural conditions. Ecology and Evolution:1–9.

Rieseberg, L. H. 1997. Hybrid origins of plant species. Annual Review of Ecology and Systematics 28:359–389.

Robertson, A. 1966. A Mathematical Model of Culling Process in Dairy Cattle. Animal Production 8:95–108.

Rockman, M. V., S. S. Skrovanek, and L. Kruglyak. 2010. Selection at Linked Sites Shapes Heritable Phenotypic Variation in *C*. elegans. Science 330:372–376.

Roda, F., L. Ambrose, G. M. Walter, H. L. L. Liu, A. Schaul, A. Lowe, P. B. Pelser et al. 2013. Genomic evidence for the parallel evolution of coastal forms in the *Senecio lautus* complex. Molecular Ecology 22:2941–2952.

Rundle, H. D., and P. Nosil. 2005. Ecological speciation. Ecology Letters 8:336–352.

Rundle, H. D., and M. C. Whitlock. 2001. A genetic interpretation of ecologically dependent isolation. Evolution 55:198–201.

Schluter, D. 2000, The Ecology of Adaptive Radiation. Oxford, Oxford University Press.

Stinchcombe, J. R., A. K. Simonsen, and M. W. Blows. 2014. Estimating Uncertainty in Multivariate Responses to Selection. Evolution 68:1188–1196.

Sweigart, A. L., and L. E. Flagel. 2015. Evidence of Natural Selection Acting on a Polymorphic Hybrid Incompatibility Locus in *Mimulus*. Genetics 199:543–554.

Tang, S., and D. C. Presgraves. 2009. Evolution of the *Drosophila* nuclear pore complex results in multiple hybrid incompatibilities. Science 323:779–782.

Walsh, B., and M. Lynch. 2018, Evolution and selection of quantitative traits. Oxford, Oxford University Press.

Walter, G. M., J. D. Aguirre, M. W. Blows, and D. Ortiz-Barrientos. 2018a. Evolution of genetic variance during adaptive radiation. The American Naturalist 191:E108–E128.

Walter, G. M., M. J. Wilkinson, J. D. Aguirre, M. W. Blows, and D. Ortiz-Barrientos. 2018b. Environmentally induced development costs underlie fitness tradeoffs. Ecology 99:1391–1401.

Walter, G. M., M. J. Wilkinson, M. E. James, T. J. Richards, J. D. Aguirre, and D. Ortiz-Barrientos. 2016. Diversification across a heterogeneous landscape. Evolution 70:1979–1992.

