## Supplementary Material for "Local adaptation and hybrid failure share a common genetic basis"

**Contents**

**Table S1:** Individuals and families used to create the ARF

**Fig S1:** Comparison of random versus observed genetic variance for eigenvectors describing GxE

**Table S2:** G-matrices for GxE at each life history stage

**Fig S2:** Comparison of random versus observed differences in selection

**Table S3:** G-matrix for F4 morphology in the glasshouse

**Supplementary methods**

**Table S1:** Numbers of genotypes used to create the ARF.

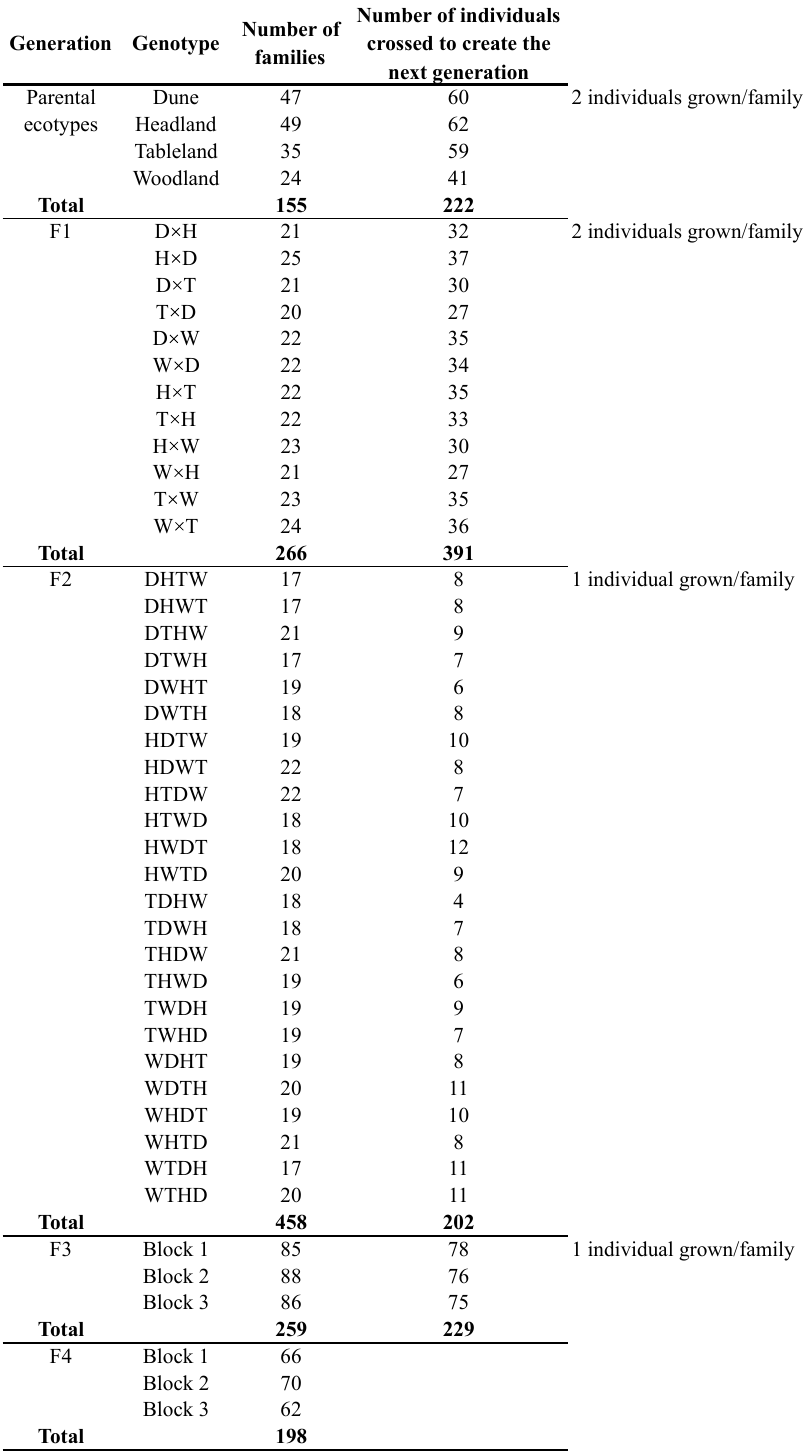

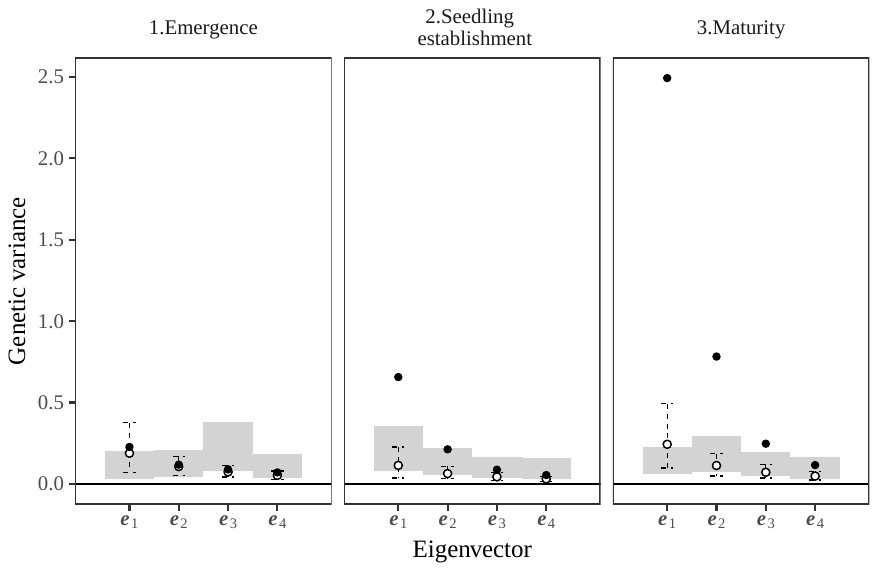

**Figure S1:** Comparing the amount of genetic variance described by eigenvectors representing the observed (filled circles) versus random matrices (unfilled circles and dashed lines), for each life history stage. Gray bars represent the amount of genetic variance in the randomized matrices described by the observed eigenvectors. Only the first three eigenvectors for maturity described more genetic variance than expected by random sampling. Credible intervals represent 95% HPD intervals.

**Table S2:** Posterior mean matrices estimating additive genetic variance in field fitness for each environment, and covariance among environments. Field fitness was analyzed separately for **A)** emergence, **B)** seedling establishment and **C)** maturity. Genetic correlations are presented below the diagonal, and covariances above the diagonal. 95% HPD intervals are presented in parentheses.

| **A)** Emergence | |  |  |  |
| --- | --- | --- | --- | --- |
|  | Dune | Headland | Tableland | Woodland |
| Dune | 0.115 (0,0.415) | 0.005 (-0.089,0.101) | 0.005 (-0.134,0.17) | 0.009 (-0.073,0.129) |
| Headland | 0.04 (-0.83,0.85) | 0.075 (0,0.259) | 0.009 (-0.108,0.136) | 0.008 (-0.054,0.099) |
| Tableland | 0.02 (-0.81,0.85) | 0.06 (-0.77,0.88) | 0.225 (0,0.691) | 0.016 (-0.108,0.151) |
| Woodland | 0.07 (-0.74,0.88) | 0.09 (-0.73,0.88) | 0.09 (-0.75,0.92) | 0.093 (0,0.358) |
| **B)** Seedling establishment | |  |  |  |
|  | Dune | Headland | Tableland | Woodland |
| Dune | 0.272 (0,0.604) | 0.091 (-0.026,0.262) | 0.071 (-0.041,0.237) | 0.154 (-0.079,0.408) |
| Headland | 0.5 (-0.27,0.98) | 0.114 (0,0.308) | 0.053 (-0.016,0.185) | 0.111 (-0.042,0.31) |
| Tableland | 0.4 (-0.45,0.97) | 0.44 (-0.42,0.99) | 0.107 (0,0.31) | 0.085 (-0.064,0.281) |
| Woodland | 0.42 (-0.2,0.94) | 0.45 (-0.28,0.98) | 0.35 (-0.39,0.95) | 0.519 (0.025,0.988) |
| **C)** Maturity | |  |  |  |
|  | Dune | Headland | Tableland | Woodland |
| Dune | 1.138 (0.371,1.956) | 0.538 (0.11,0.965) | 0.662 (0.221,1.237) | 0.419 (-0.356,1.247) |
| Headland | 0.8 (0.54,0.99) | 0.42 (0.086,0.837) | 0.433 (0.12,0.811) | 0.311 (-0.193,0.879) |
| Tableland | 0.68 (0.34,0.99) | 0.73 (0.34,0.98) | 0.894 (0.348,1.54) | 0.65 (-0.029,1.561) |
| Woodland | 0.42 (-0.29,0.99) | 0.5 (-0.23,0.99) | 0.66 (0.07,0.99) | 1.186 (0,3.095) |

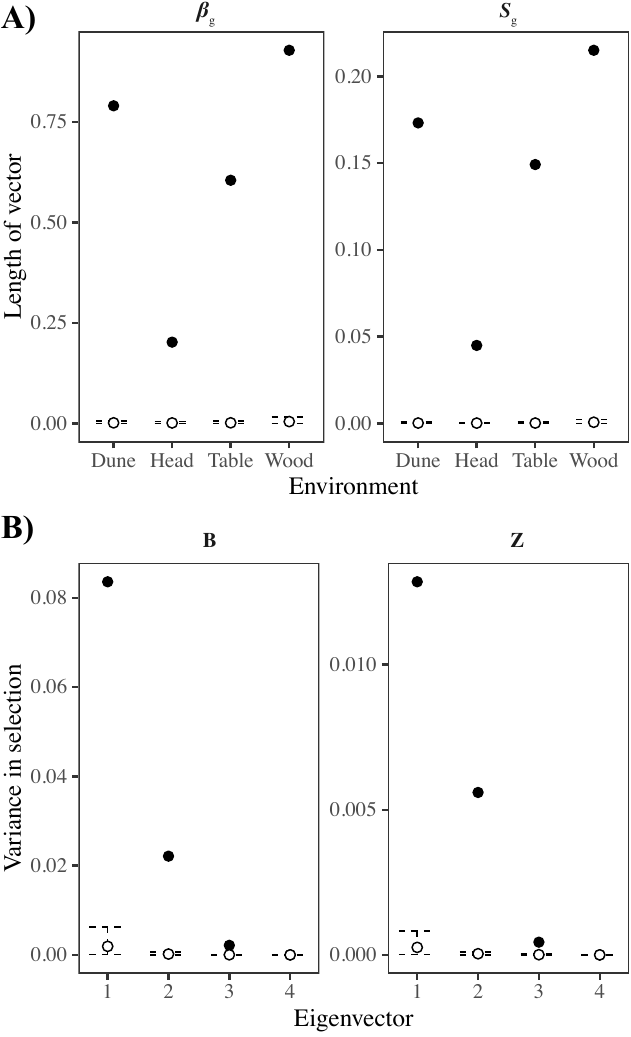

**Fig S2: A)** Comparing the length of ***β*** and ***s***_g_ in each environment, for the observed versus and random distribution. All observed vectors are greater than expected under random sampling. **B)** Comparing the amount of variance in **B** and **Z**, for the observed versus random distribution. Differences in ***β*** and ***s***_g_ were greater than expected under random sampling. Credible intervals represent 95% HPD intervals.

**Table S3:** G-matrix estimated on the morphology data from experiment 2. Numbers in bold show the traits that were chosen for the Robertson-Price Identity as they possessed a heritability greater than 0.1. Covariances are presented above the diagonal, with correlations below the diagonal. 95% HPD intervals are presented in parentheses.

|  | Height | Area | P2A2 | Circularity | Nindents | Indent width | Indent depth |
| --- | --- | --- | --- | --- | --- | --- | --- |
| Height | **0.435**  **(0,0.896)** | 0.079  (-0.091,0.277) | -0.064  (-0.261,0.144) | 0.008  (-0.098,0.114) | -0.029  (-0.155,0.063) | 0.089  (-0.095,0.266) | -0.038  (-0.182,0.052) |
| Area | 0.23  (-0.36,0.75) | **0.283**  **(0,0.567)** | -0.167  (-0.385,0.031) | 0.024  (-0.061,0.176) | -0.022  (-0.154,0.091) | 0.142  (-0.029,0.335) | -0.057  (-0.223,0.035) |
| P2A2 | -0.21  (-0.87,0.37) | -0.55  (-0.95,0.02) | **0.295**  **(0,0.586)** | -0.047  (-0.232,0.043) | 0.013  (-0.114,0.141) | -0.237  (-0.445,0.003) | 0.079  (-0.043,0.27) |
| Circularity | 0.07  (-0.64,0.75) | 0.17  (-0.68,0.9) | -0.3  (-0.98,0.7) | 0.047  (0,0.179) | 0.008  (-0.046,0.065) | 0.021  (-0.11,0.134) | -0.021  (-0.114,0.029) |
| Nindents | -0.18  (-0.83,0.6) | -0.16  (-0.85,0.75) | 0.15  (-0.84,0.94) | 0.06  (-0.76,0.95) | 0.052  (0,0.167) | -0.052  (-0.218,0.079) | -0.01  (-0.086,0.037) |
| Indent width | 0.26  (-0.28,0.84) | 0.47  (-0.06,0.94) | -0.75  (-0.99,-0.37) | 0.19  (-0.74,0.96) | -0.29  (-0.94,0.77) | **0.342**  **(0.06,0.661)** | -0.057  (-0.184,0.056) |
| Indent depth | -0.22  (-0.9,0.44) | -0.35  (-0.96,0.51) | 0.45  (-0.48,0.97) | -0.24  (-0.95,0.6) | -0.04  (-0.88,0.8) | -0.4  (-0.96,0.54) | 0.071  (0,0.216) |

**Supplementary methods**

**Implementation of Bayesian models.** For each analysis, we implemented Markov chains of different lengths (listed in Table S5), while ensuring that we included a sufficient burn-in period and thinning interval to sample the parameters with autocorrelation values of less than 0.05 and effective sample sizes exceeding 85% of the total number of samples, for all parameters. We used uninformative parameter expanded priors and checked their sensitivity by re-implementing all models while adjusting the parameters and ensuring the posterior distribution did not change.

**Table S5:** Number of MCMC iterations used to implement each analysis. NA = Not Applicable.

| **Analysis** | **Equation** | **Thinning interval** | **Burn-in period** | **Total iterations** | **Reduction of total MCMC iterations for randomized data** |
| --- | --- | --- | --- | --- | --- |
| Estimate of mean fitness for F4 and parental ecotypes | Equation 1 | 5,000 | 100,000 | 5,100,000 | NA |
| Estimate genetic variance for F4 fitness | Equation 2 | 2,000 | 100,000 | 2,100,000 | 200 |
| G-matrix to choose phenotypic traits for equation 4 (Robertson-Price Identity) | Equation 3 | 5,000 | 500,000 | 5,500,000 | NA |
| Robertson-Price Identity (selection on phenotype) | Equation 4 | 2,500 | 100,000 | 2,600,000 | 165 |

**Randomizations of observed data.** For the analyses estimating genetic variance, comparing estimates of genetic variance with zero provides an uninformative test of significance because estimates are restricted to be greater than zero (positive-definite). To create an informative significance test, we re-implemented each model with randomized data, created by shuffling the parental information. For each model implemented on the observed data, we re-implemented the same model on 1,000 randomizations of the data, and extracted the posterior mean for each randomization. We then compared the distribution of means from models conducted on the randomizations, to the mean of the observed posterior distribution. If the mean of the observed distribution occurred outside the 95% Highest Posterior Density (HPD) interval for the random distribution, we took this as evidence that we captured biologically important information for the comparison of interest. As we were only interested in estimating the posterior mean of models implemented on each randomization of the data, we could reduce computing time by reducing the total number of sampling iterations. To do so, we maintained the same burn-in period and sampling interval to ensure an identical mixing of MCMC chains, reducing only the total number of sampling iterations to the number required to obtain a stable estimate of the mean. We calculated the number of sampling iterations required using the models implemented on the observed data, which was specific to each analysis (Table S5).
